# Conditioned medium from painful schwannomatosis tumors increases pain behaviors in mice

**DOI:** 10.1101/2024.04.06.588413

**Authors:** Randy Rubright, Michael J. Caterina, Allan Belzberg, Kimberly Laskie Ostrow

## Abstract

The majority of schwannomatosis (SWN) patients experience debilitating pain. Yet, it is not known why only some schwannomas cause pain or whether mutations in SWN-related genes, (SMARCB1 or LZTR1) differentially influence pain signaling pathways. Here, we established cell lines from SWN tumors resected from patients with varying degrees of pain and bearing mutations in different SWN-related mutations. Compared with conditioned medium (CM) collected from “nonpainful” SWN tumors, CM from “painful” SWN tumors contained elevated levels of specific inflammatory cytokines (IL-6, IL-8, VEGF), and was able to enhance sensory neuron responsiveness to noxious TRPV1 and TRPA1 agonists in vitro. In in vivo studies, injection of CM from painful SWN into the hind paws of healthy mice evoked both more acute pain behavior and greater enhancement of mechanical stimulus-evoked behavioral responses than did CM from nonpainful SWN. Furthermore, the behavioral effects of painful CM differed as a function of the SWN-related gene mutations identified in the tumors of origin. Painful SMARCB1 mutant CM, for example, sensitized mice to mechanical stimulation at low forces, compared to non-painful tumor CM and control media, but this effect waned over time. In contrast, CM from a painful tumor with no detectable mutation in either SMARCB1 or LZTR1 caused the greatest increase in responsiveness to low mechanical forces and this effect lasted for 2 days post-injection. These experiments establish a paradigm for examining the mechanisms by which painful SWN tumors bearing different mutations produce their sensory effects and will thus facilitate better understanding and, potentially, treatment of the pain endured by SWN patients.

## Introduction

Patients with schwannomatosis (SWN) develop multiple tumors along major peripheral nerves and often experience extreme pain[1]. In many cases, patients experience pain prior to the detection of a palpable mass. Pain may or may not be directly related to the size or location of the tumor, and not all tumors are painful [2]. Surgical resection is the current standard of care for patients who harbor painful schwannomas, however tumor removal may not provide lasting pain relief due to tumor growth and is complicated by overall tumor burden [3]. Many patients have been trialed on up to 10 different medications for pain with little relief [4]. We do not understand the etiology of schwannomatosis related pain. Neuropathic, nociceptive, and inflammatory pain have all been described by patients [5] and different symptoms of pain have been described even within a patient [6].

To complicate matters, schwannomatosis as a disease entity has recently been divided into subgroups based on gene mutations found in patients with multiple schwannomas [7]. Mutations in either of two genes on chromosome 22q11 (LZTR1 or SMARCB1) account for ∼90% of familial schwannomatosis. In sporadic disease, these mutations account for ∼45 % of cases. Patients harbor mutations in either SMARCB1 or LZTR1 but not both. Schwannomatosis is therefore classified as 1) SMARCB1 related, 2) LZTR1 related, or 3) Schwannomatosis Not Otherwise Specified (NOS)/ Not Elsewhere Classified (NEC). The NOS/NEC group does not harbor a mutation in either SMARCB1 or LZTR1. It is currently unknown whether mutation status influences the painful phenotype of schwannomatosis, although it has been suggested that patients with mutations in LZTR1 express increased incidence of pain [8].

Due to the heterogeneity of the painful phenotype seen between and within schwannomatosis patients, we speculate that there is an alteration occurring in the schwannoma itself and that this alters the behavior of nociceptive neurons. We are particularly interested in how the tumors’ “secretome” affects sensory neurons. To test our hypothesis, we established immortalized (SV40 large T antigen-transfected) cell lines from resected tumors from SWN patients. Tumors were selected based on varying degrees of pain (no, mild, or severe pain). Cell lines demonstrated the same gene expression as the tumors from which they were derived, as confirmed by Illumina HT-12 microarray expression analysis [9]. In our previous study, we found that conditioned medium (CM) collected from painful SWN tumors, but not that from nonpainful SWN tumors, contained increased amounts of multiple cytokines, upregulated the expression of pain-associated genes in cultured dorsal root ganglion (DRG) neurons, and increased the responsiveness of these neurons to noxious agonists for transient receptor potential vanilloid 1 (TRPV1) and transient receptor potential ankyrin 1 (TRPA1) channels in vitro [10].

Our prior in vitro studies showed convincing evidence that substances secreted by SWN tumors sensitize neurons and alter neuronal gene expression. However, they did not address whether this might translate into pain alterations in vivo. In this study, using well-established methods to quantify pain levels in healthy mice, we therefore assessed the effects of SWN CM on acute and mechanically-evoked pain behaviors. In addition, we examined the influence of SWN mutations on cytokine secretion in tumor CM and on the sensitivity of CM-treated sensory neurons to TRPA1 and TRPV1 agonists. This study expands our previously published in vitro findings and validates the notion of mechanistic heterogeneity in the pain experienced in schwannomatosis.

## Results

### Painful SWN tumor CM contains elevated levels of cytokines and chemokines

In our prior study [10], we published a qualitative description of elevated levels of cytokines in CM from painful (n=4) and non-painful (n=3) tumors using the Proteome Profiler Human Cytokine XL Array (R&D Systems). To validate and further explore these differences, we narrowed down our analysis to 9 candidates (CCL2, IL-6, IL-8, VEGF, GDF-15, CXCL1, CXCL5, CCL20 and GM-CSF) indicated by the cytokine array to be highly expressed and performed quantitative ELISAs on an expanded cohort of SWN tumor CMs (Painful CM n=12 samples; Non-painful CM n=8 samples) **(Figure 1)**. These assays revealed significantly higher levels of IL-6 *(p=0*.*03)*, IL-8 *(p=0*.*01)*, VEGF *(p=0*.*008)* and GDF-15 *(p=0*.*01)* in painful CM, compared with non-painful CM **(Figure 1)**. These findings both support and extend our observation of elevated cytokine release by painful SWN tumors. As many of these cytokines have been previously linked to pain, they also provide a plausible mechanism for nociceptor sensitization by painful SWN CM.

**Figure 1:**
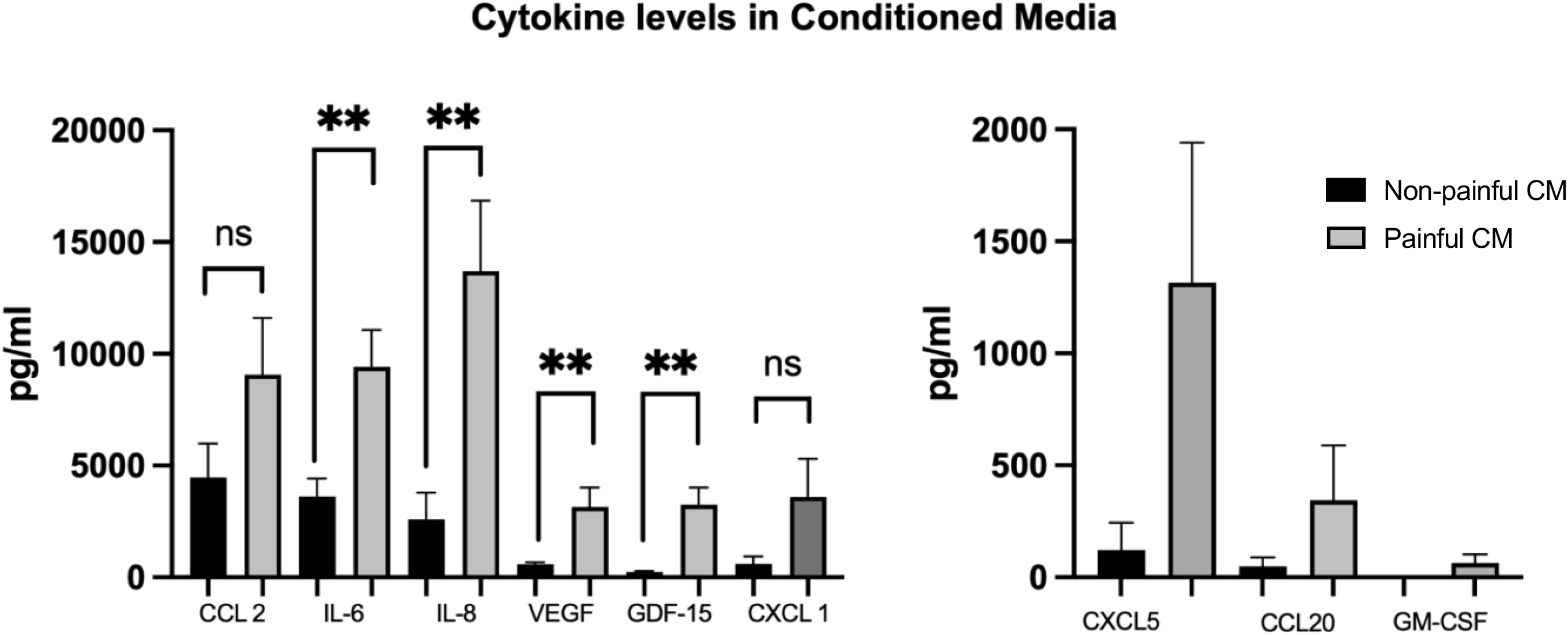
Painful Schwannoma CM contains elevated levels of secreted cytokines and chemokines. Painful CM (n=12 samples, gray bars) and non-painful CM (n=8 samples, black bars) were examined by ELISA to measure amounts of specific cytokines. CCL2, IL-6, IL-8, VEGF, GDF-15, CXCL1, CXCl5, CCl20 and GM-CSF were tested. Painful CM contains significantly higher levels of IL-6 *(p=0*.*03)*, IL-8 *(p=0*.*01)*, VEGF *(p=0*.*008)* and GDF-15 *(p=0*.*01)*. Unpaired t-test with Welch’s correction was used to determine significance.

### Conditioned media from painful schwannomas causes an increase in pain behaviors in mice

Our prior analysis of SWN CM effects was confined to in vitro assays [10]. To assess the effects of SWN CM in vivo, we therefore injected one hindpaw of healthy C57Bl6 mice with conditioned media from painful (n=40 mice) or non-painful tumors (n=30 mice), or with non-conditioned medium (Dulbecco’s modified eagle’s medium/10% FBS/2uM forskolin; n=40 mice). We utilized media from 4 separate painful tumors, and 3 separate non-painful tumors that had been analyzed in our prior in vitro study. To test for acute pain responses to CM, mice were video recorded for 10 minutes following injection. Mice that received injections of CM from non-painful tumors showed increased paw licking or flinching, compared to those injected with control media (p=0.0002). Painful CM caused mice to exhibit an even greater acute behavioral response compared to control media (p<0.0001) and non-painful tumor CM (p=0.01) **(Figure 2a**).

**Figure 2:**
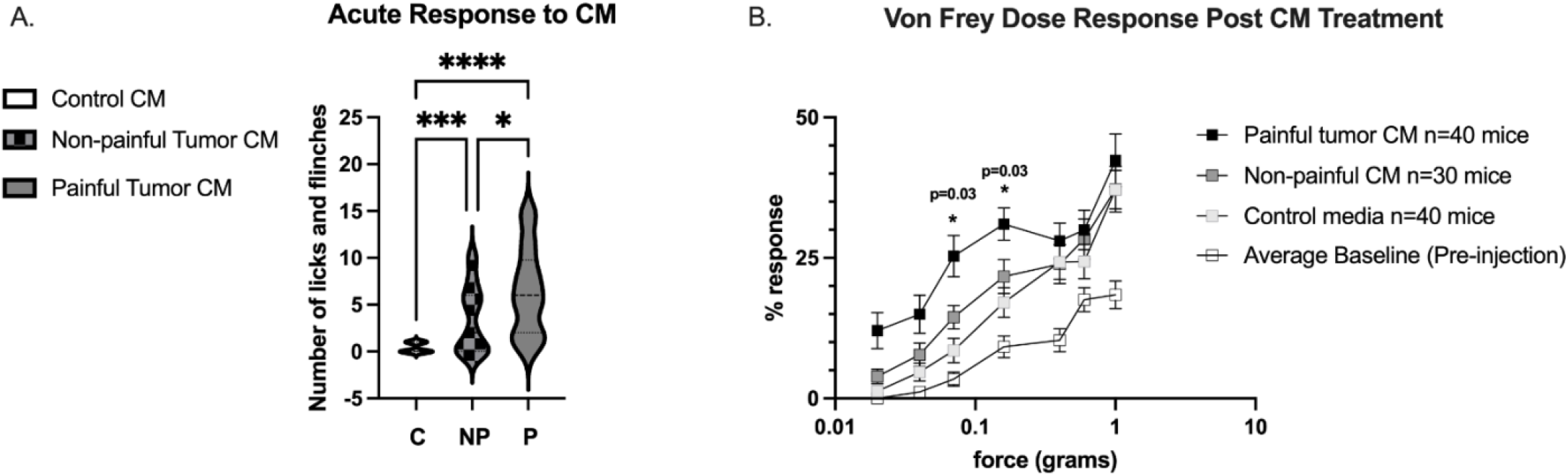
Conditioned media from painful SWN tumors cause increased pain behaviors upon intra-plantar paw injection. Ten microliters of CM were injected into the hindpaw of C57black6 mice. CM from 4 painful tumors n=40 mice and CM from 3 non-painful tumors n=30 mice were tested. Control media (non-conditioned Schwann cell media, DMEM 10% FBS, and 2% forskolin) was injected into an additional 40 mice. ***a). Acute pain response*** was assessed by examining the number of times a mouse licked or flinched the affected paw. CM from painful tumors significantly increased the number of licks/flinches over a 10-minute observation period compared to non-painful CM (p=0.01) and control media (p<0.0001). Brown-Forsythe ANOVA test with Dunnett’s T3 multiple comparisons was used to determine significance ***b). Response to evoked mechanical pain was examined using Von Frey filaments***. Increasing forces from 0.02g to 1g were tested. Each filament was tested 6 times. The percent response was calculated for each mouse at each force. Response was measured prior to injection (open squares) and one hour after the final injection. Painful CM induced an increased response to light touch (black squares) compared to control media (light gray squares) at forces less than 0.4g. Painful media caused increased response compared to non-painful CM (dark gray squares) at 0.16 g *(p=0*.*03)* and 0.07g *(p=0*.*03)*. Two-way ANOVA with repeated measures, and Tukey’s multiple comparisons were used to determine significance between groups at different forces

Following the observation period, and to simulate our in vitro CM sensitization study, mice were injected once daily with CM for two additional days. One hour after the final injection, the mice were examined for mechanical sensitivity using the von Frey filament assay. Pre-injection, mice exhibited the expected pattern of increasing response frequency with increasing stimulus force. Following injection, painful CM treated mice exhibited hypersensitivity to lower forces (0.02g, 0.04g, 0.07g, and 0.16g), compared with those injected with control media or non-painful CM **(Figure 2b; Table 1; Supplementary figure 1)**.

**Table 1.**
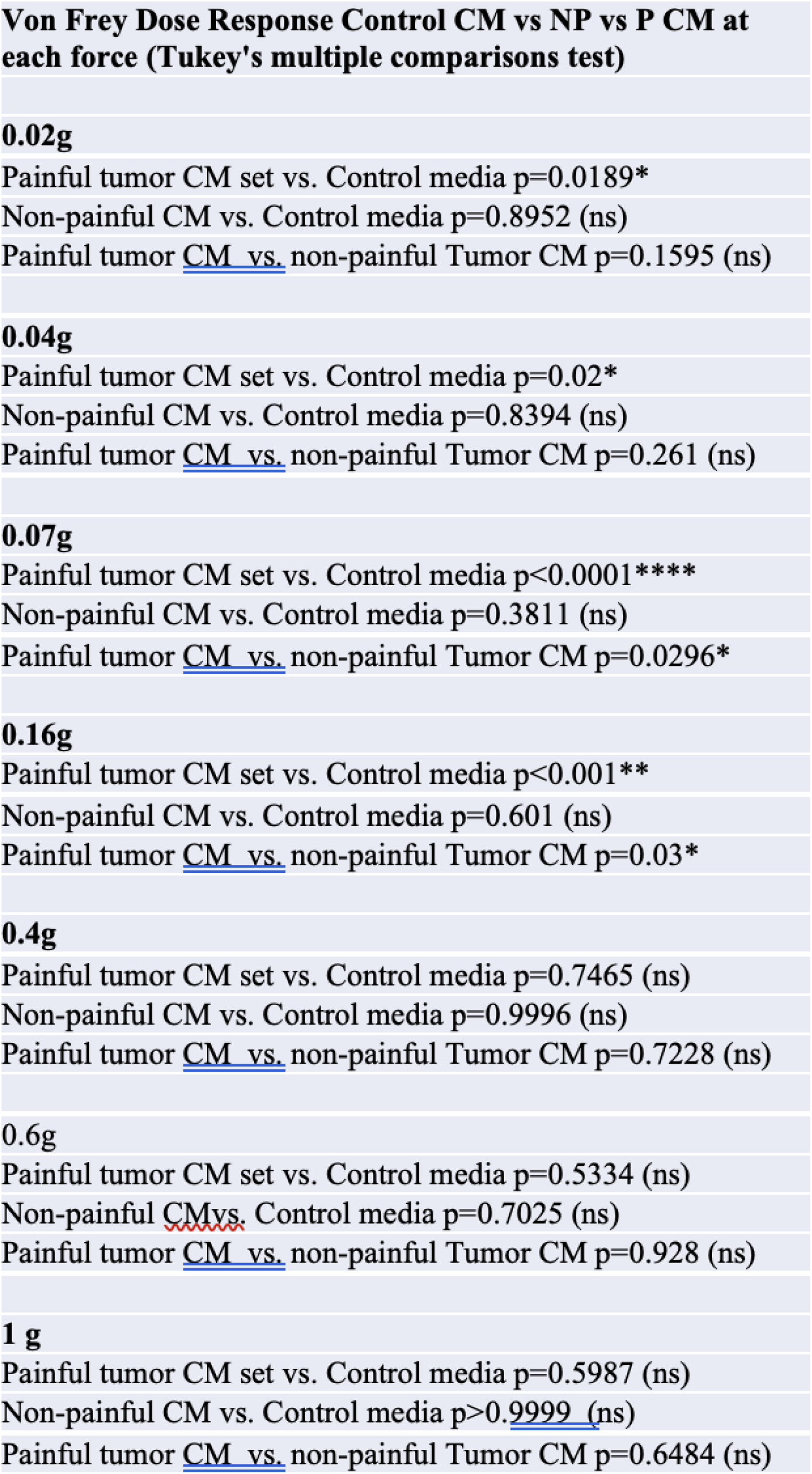

### In vitro and in vivo response to painful and non-painful CM segregated by mutation group

#### Effect of SMARCB1, LZTR1, and NOS mutant tumor CM on cultured DRG neurons

Our previous work demonstrated that cultured mouse dorsal root ganglion (DRG) neurons that were pretreated with CM from painful tumors, but not that from non-painful tumors, were hypersensitized to low doses of capsaicin (∼25nM), a TRPV1 agonist and/or cinnamaldehyde (∼200uM), a TRPA1 agonist. We also noted that each painful tumor tested had a slightly different effect on the cultured neurons [10]. At the time of that publication, mutations were not considered as diagnostic criteria for SWN and were thus not examined. We subsequently collected painful (patient-reported pain score > 7) and non-painful (patient-reported pain score < 3) tumors with known mutations in SMARCB1 and LZTR1. We also made use of confirmed SWN NOS tumors where sequencing of SMARCB1 and LZTR1 detected no mutations in either gene. We created cell lines and collected CM from these tumors. We then repeated our sensory neuron sensitization experiments using these additional CMs from pain score- and mutation-matched groups. Mouse DRGs were dissociated and the resulting cells were plated and incubated with CM for 48 hours. Following incubation, the neurons were subjected to fluorescent calcium imaging analysis during challenge with low-dose capsaicin and cinnamaldehyde as previously described [10]. Low-dose capsaicin caused a rapid influx of calcium in a subset of neurons that quickly returned to baseline levels upon washout, resulting in a bell-shaped curve, while low-dose cinnamaldehyde evoked a slower increase in calcium in a subset of neurons that very slowly declined upon washout but did not return to baseline levels, resulting in an overall larger area under the curve **(Figure 3a)**. All painful CMs sensitized DRG neurons to low dose capsaicin compared with their matched non-painful counterparts as demonstrated by either an increased magnitude of response (among responsive cells) or an increase in the percentage of cells responding to capsaicin **(Figure 3b, Table 2)**. However, differences were seen between painful CM groups based on mutation. Painful SMARCB1 mut CM (AUC=16.2) and LZTR1 mut CM (AUC 15.6) enhanced the response to low-dose capsaicin more than NOS mut CM (AUC=11) (*p=0*.*0001; p=0*.*003*, respectively) **(Figure 3b)**. Conversely, painful NOS mut CM enhanced the response to low-dose cinnamaldehyde (AUC 21.3) more than non-painful NOS tumor CM (AUC=11.8) *(p=0*.*01)* or painful LZTR1 tumor CM (AUC=9.59) *(p=0*.*007)* **(Figure 3c)**. While pretreatment of DRG cells with painful SMARCB1 CM and non-painful SMARCB1 CM both increased the magnitude of response to cinnamaldehyde, compared to control CM, painful SMARCB1 boosted the percentage of cells sensitized to cinnamaldehyde (44% painful SMARCB1 CM vs 17% non-painful SMARCB1 CM *p=0*.*0001*) (**Figure 3c, Table 2**).

**Table 2:**

| **Number of cells responsive to TRP agonists** |  |  |
| --- | --- | --- |
|  | **25nM Capsacin** | **200uM Cinnamaldehyde** |
| **Control CM** | 56/230 (24%) | 22/147 (15%) |
| **NP LZTR1** | 28/148 (19%) | 44/207 (21%) |
| **P LZTR1** | 293/540 (54%)**** | 48/209 (24%) |
| **NP SMARCB1** | 106/418 (25%) | 25/149 (17%) |
| **P SMARCB1** | 395/597 (66%)**** | 46/104 (44%)**** |
| **NP NOS** | 61/226 (27%) | 86/316 (27%) |
| **P NOS** | 151/319 (47%)**** | 226/414 (54%)**** |
|  | **** p<0.0001 |  |

Chi-square analysis p<0.05 is significant

**Figure 3:**
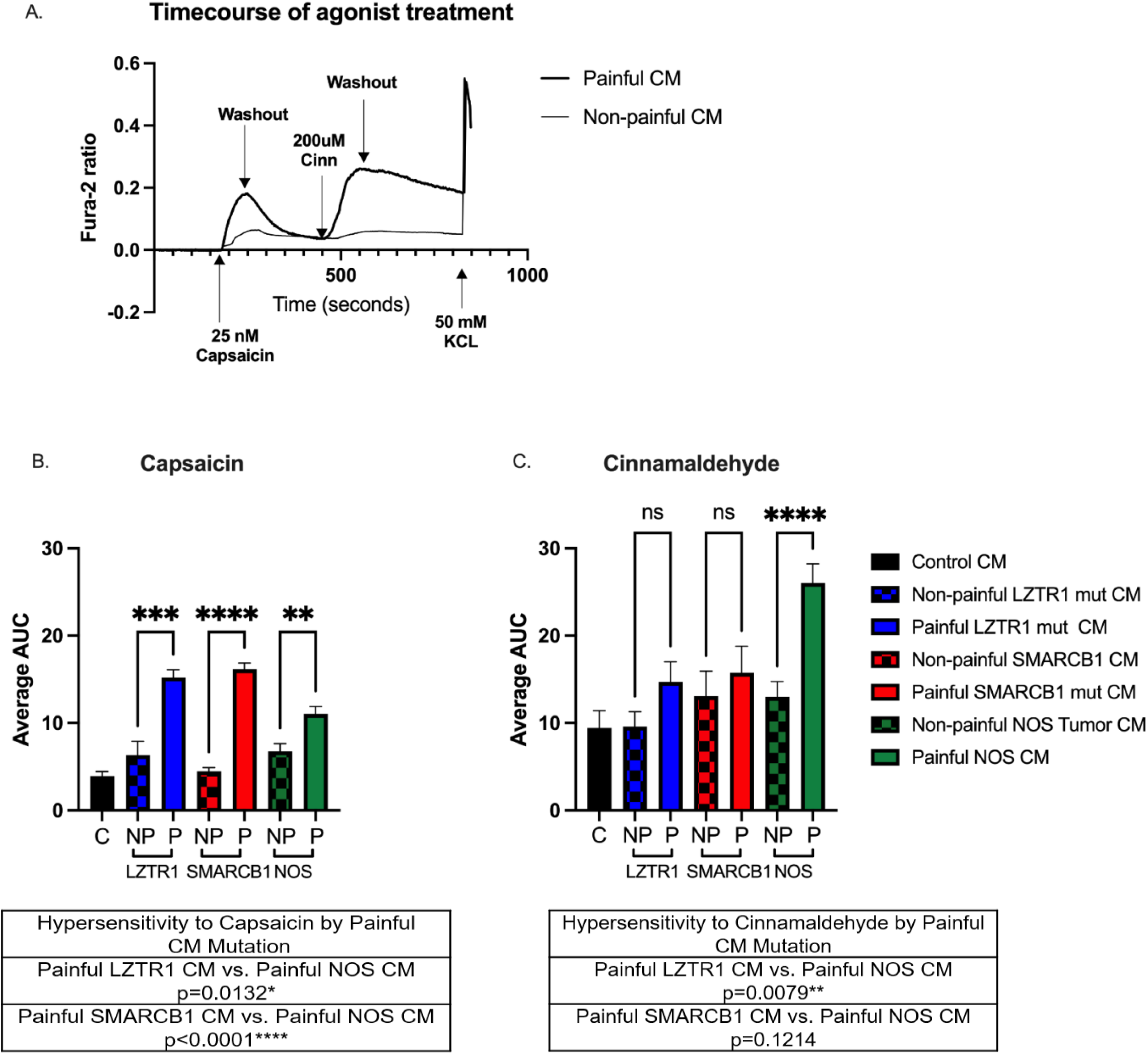
Conditioned media from painful SWN tumors sensitize DRG neurons to the TRPV1 agonist, capsaicin, and the TPRA1 agonist, cinnamaldehyde. ***a). Representative calcium imaging trace*** DRG cells were pre-treated with painful schwannoma CM, non-painful schwannoma CM, or control media for 48 hours before imaging. Fura-2 ratio measurements were recorded at 2 second intervals. The datapoints were corrected for differing baseline readings of Fura-2. Area under the curve was calculated for each treatment. Six coverslips of DRGs were tested per CM. *Brown-Forsythe ANOVA test* with Dunnett’s T3 multiple comparisons was employed to determine statistical significance p<0.05 is considered significant ***b) TRPV1 agonist, Capsaicin stimulus*** CM from painful tumors increased the responsiveness of the DRGs to capsaicin as calculated by area under the curve. Solid filled bars are painful CM. Checkered bars are non-painful CM. Comparisons between painful CMs of different mutations were also assessed. ***c) TRPA1 agonist, Cinnamaldehyde stimulus*** CM from painful NOS/NEC tumor sensitized DRG neurons to low dose cinnamaldehyde *(p=0*.*02)*. Additionally, comparisons between painful CMs of different mutations were also examined.

#### Acute Behavioral Response to CM differs by mutation

The mouse behavior studies described in Figure 2 were next repeated, considering mutation status, using CM from our expanded cohort of samples. The CM examined in Figure 3 was injected into the foot pad of C57Bl6 mice (10/group). Mice were observed for 10 minutes and the number of times a mouse licked or flinched the injected paw was recorded. All painful SWN CMs caused an increase in licking and flinching compared to control media **(Figure 4)**. However, only LZTR1 painful CM caused an increased acute pain response compared to their non-painful LZTR1 CM counterpart *(p=0*.*006)*. Further comparison between tumor genotypes revealed that CM from painful tumors with LZTR1 mutations caused an increase in acute pain response compared to painful CM from SMARCB1 (*p=0*.*0002*) and NOS tumors (*p=0*.*0001*) **(Figure 4)**. In addition, non-painful LZTR1 CM also elicited more of an acute pain response than control media *(p=0*.*0001)*, non-painful SMARCB1 *(p=0*.*0007)* and non-painful NOS CM *(p=0*.*0001)*. No significant differences in cumulative licks and flinches were noted between painful and non-painful tumor CM from SMARCB1 or NOS tumors.

**Figure 4:**
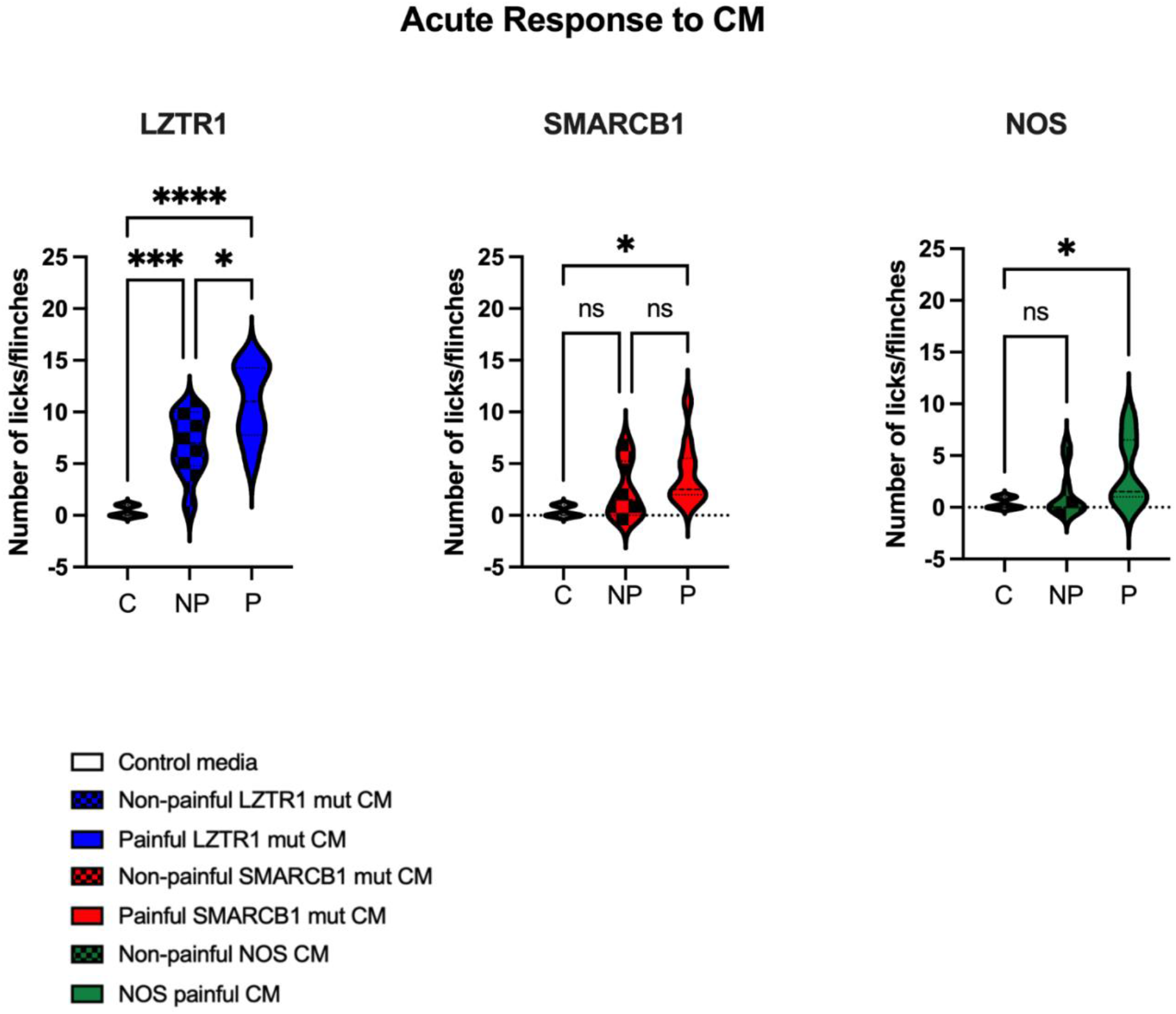
Acute Response to CM differs by mutation. Painful CM from a SMARCB1 mut tumor, LZTR1 mut tumor, and NOS mut tumor were tested. All painful tumors had a pain score greater than 7. One non-painful tumor (pain score <3) from each mutation group was used for comparison. Ten mice were injected per CM. All painful SWN CMs caused an increase in licking and flinching compared to control media. LZTR1 painful CM caused an increased acute pain response compared to non-painful LZTR1 CM compared to control media *(p=0*.*006, p=0*.*0001, respectively). Brown-Forsythe* ANOVA with Dunnett’s T3 multiple comparisons tests were used to determine significance.

### CM-evoked behavioral hypersensitivity to mechanical stimuli varies by mutation

We next examined the responsiveness of CM-injected mice to punctate mechanical stimuli evoked by von Frey monofilaments (0.02 to 0.07g). Mice were assayed one hour and again 24 and 48 hours after the final CM injection. CM from painful SMARCB1mutant tumor sensitized the injected paw to evoked mechanical pain, compared to non-painful SMARCB1 CM at low forces (0.04g *p=0*.*03*; 0.07g *p=0*.*01*) **(Figure 5a)**. This effect subsided over the ensuing 24 hours **(Figure 5b)**. Forty-eight hours post-injection, no differences between painful and non-painful CM were detected **(Figure 5c)**. An even more prominent sensitization to mechanical stimuli was produced by CM from painful NOS tumor. Injection of this CM elicited a greater degree of mechanical responsiveness than that from nonpainful NOS tumor at the lowest force tested (0.02g) (NOS mut painful vs NOS mut non-painful *p=0*.*0016*) as well as at 0.04g (NOS mut painful vs NOS mut non-painful *p<0*.*0001)* and 0.07g *(p<0*.*0001)* **(Figure 5a)**. Moreover, this effect was durable, in that hypersensitivity persisted 24 hours post-injection (0.04g *p=0*.*0017*; 0.07g *p=0*.*0002*) **(Figure 5b**). Forty-eight hours post-injection, mice treated with painful NOS CM still demonstrated an apparently increased response to light touch, but the difference from non-painful NOS CM was not statistically significant **(Figure 5c)**. No statistically significant differences in punctate mechanical sensitivity were observed between non-painful LZTR1-related CM and painful LZTR1-related CM at any time point post injection **(Figures 5 a,b,c)**.

**Figure 5.**
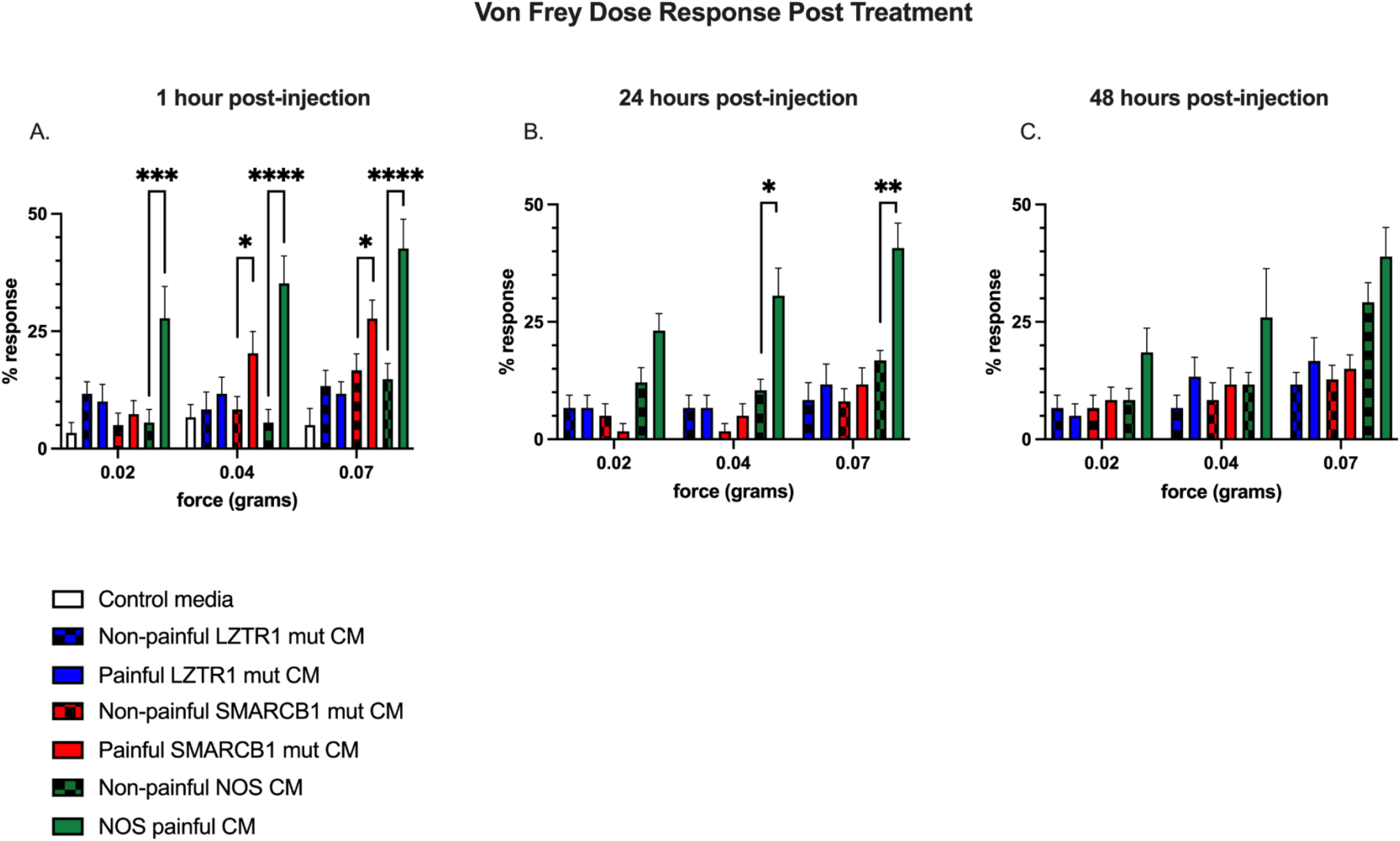
Response to painful stimuli differs between SWN mutational groups. Evoked mechanical hypersensitivity to light touch (0.02g, 0.04g, and 0.07g) was assayed using Von Frey (VF) filaments. Percent response was calculated as in Figure 2b. Four separate rounds of VF were performed on each mouse over the course of the experiment. The first round was performed prior to any injections with the subsequent rounds being performed 1 hour following the final injection, 24 hours post final injection, and 48 hours post final injection. **5a)**. No differences were seen between mutation groups of non-painful CM (checked bars). Painful NOS mut CM (Green Bars) and SMARCB1 mut CM (Red Bars) caused an increase in response to light touch at 0.04g and 0.07g compared to corresponding non-painful CM (checked bars), while painful LZTR1 (Blue bars) did not affect response to light touch at any force. Painful NOS mut CM gave a significant response to light touch at the lowest Von Frey filament force tested (0.02g, *p=0*.*01*). **5b)**. Twenty-four hours post-injection, painful NOS CM maintained a significant response to light touch compared to non-painful NOS CM at 0.04g *(p=0*.*02)* and 0.07g *(p=0*.*004)*. **5c)**. Forty-eight hours post-injection, increased hypersensitivity to light touch was still demonstrated in the painful NOS CM treated mouse cohort. Two-way ANOVA with repeated measures, and Tukey’s multiple comparisons were used to determine significance.

### Levels of specific cytokines and chemokines in the injection cohort CMs

Finally, we assayed the CM used in the in vivo experiments above for levels of CCL2, IL-6, IL-8, VEGF, GDF-15, CXCL1, CXCL5, CCL20 and GM-CSF, using ELISA assays (R&D systems). Levels of IL-6, IL-8, and VEGF were higher in painful CM than non-painful CM regardless of mutation group (Figure 1). When comparing painful CMs by mutation, painful NOS CM **(Figure 6, Green bars**) contained higher levels of IL-8, CCL2, and CCL20 than Painful SMARCB1CM **(Figure 6, Red bars**) and Painful LZTR1 CM (**Figure 6, Blue bars**). Painful LZTR1 CM contained higher levels of GDF-15, CXCL1 and GM-CSF than Painful NOS and Painful SMARCB1 CM.

**Figure 6:**
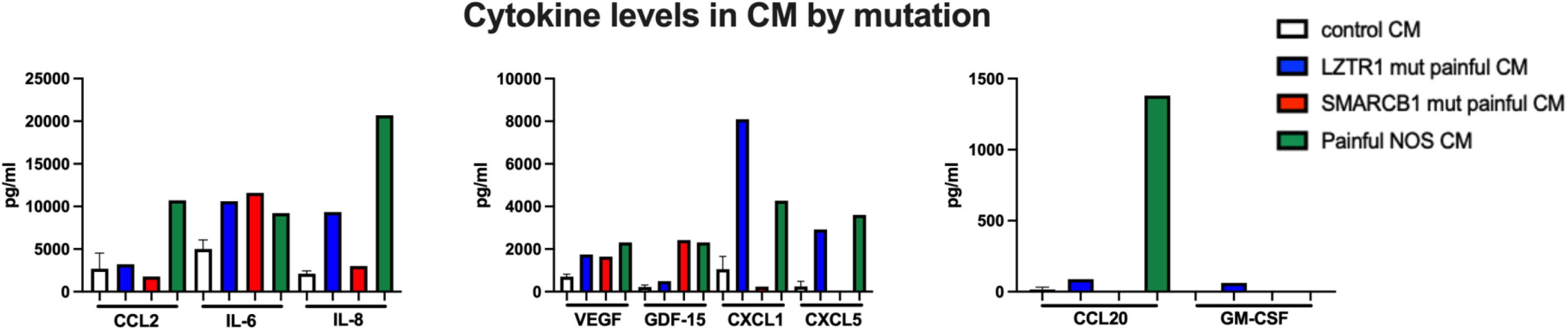
Cytokine levels in CM by mutation status. ELISA was used to examine the levels of our candidate panel of cytokines/chemokines in painful CM from our injection cohort. CCL2, IL-6, IL-8, VEGF, GDF-15, and CXCL1, CXCL5, CCL20 and GM-CSF were tested. Painful NOS CM (Green Bars) contains higher levels of IL-8, CCL2, and CCL20 than Painful SMARCB1CM (red bars) and Painful LZTR1 CM (Blue bars). Painful LZTR1 contained higher levels of GDF-15, CXCL1 and GM-CSF than Painful NOS and Painful SMARCB1 CM.

## Discussion

Patients with schwannomatosis describe their pain experience with great diversity. While the majority of patients experience chronic pain, symptoms can be localized to a tumor or may be diffuse. In some cases a particular tumor may not cause any pain. Our prior study focused on deciphering the molecular mechanisms of painful tumors (Pain score >7) compared to “non-painful” tumors (pain score <3) [10]. Those in vitro experiments revealed that CM collected from painful SWN tumors, but not that from nonpainful SWN tumors, contained increased amounts of multiple cytokines, upregulated the expression of pain-associated genes in sensory neuron cultures, and increased neuronal responses to noxious TRPV1 and TRPA1 agonists [10]. In the present study, these and additional CM samples were injected in vivo to assess whether the secretome of SWN cells can induce painful behaviors in mice. We found that painful tumor-derived CMs increase acute pain behaviors when injected into the glabrous skin of C57Bl6 mice. An increase in licking and flinching occurred after a single injection of painful CM (Figure 1a). Behavioral responsiveness to punctate mechanical stimuli also increased in mice injected with painful CM. While this increase between the grouped cohorts was statistically significant overall (Figure 1b), variability was evident between painful tumors (supplementary figure 1). This variability may reflect that seen with these CM samples in vitro [10]. CM samples that hypersensitized neurons to capsaicin in vitro also hypersensitized mice to punctate stimuli. Moreover, the samples that sensitized neurons to both cinnamaldehyde and capsaicin *in vitro* [10], produced the largest enhancement of mechanically-evoked pain *in vivo* (painful Tumor 3; Supplementary figures 1 & 2).

In 2022, experts in the field of peripheral nerve sheath tumors established new criteria for diagnosis of schwannomatosis [7]. All tumors of Schwann cell origin were to be classified as schwannomatosis. Neurofibromatosis type 2 was renamed to *NF2-related schwannomatosis [7, 11]*. The multiple tumor disorder we have been studying, formerly called SWN, was renamed and subdivided to reflect tumor mutation status. As a group, this disease is termed *non-NF2 schwannomatosis (non-NF2-SWN)*. Subgroups of tumors are classified as SMARCB1-related SWN, LZTR1-related SWN, or SWN not otherwise specified. Based on this reclassification, in the present study we repeated our in vitro and in vivo studies using an additional set of samples with known germline mutations in these genes. It is not known whether mutations in SMARCB1 or LZTR1 correlate with a patient’s experience with pain. However, based on our new data, we believe tumor mutation status may contribute to variability in cytokine secretion and neuronal sensitization in vitro and in vivo.

CM from painful tumors with mutations in SMARCB1 produced significant sensitization to punctate mechanical stimuli that was evident one hour after injection but subsided by 24 hours. Painful SMARCB1 CM contains elevated levels of Il-6, VEGF, and GDF-15, compared to non-painful tumor CM. Previous studies demonstrated that IL-6 induced mechanical hypersensitivity in otherwise naive rats in vivo and increased DRG neuron sensitivity to capsaicin in vitro[12]. Another study demonstrated that IL-6 injection also increased hyperalgesia in the rat footpad. Intriguingly, this effect peaked 2-3 hours after injection, was still demonstrated 6 hours post injection, but subsided 24 hours post injection [13], consistent with the kinetic profile exhibited by painful SMARCB1 CM in the present study.

Injection of painful NOS mutant CM caused an amplified, long-lasting response to punctate mechanical stimuli, even increasing the response to the lowest force tested. Painful NOS CM also contains elevated levels of IL-6 VEGF, CXCL-1, and GDF-15, compared to non-painful CM. However, it additionally contains elevated levels of IL-8, CCL2, CXCL-5, and CCL20, compared to painful CM from SMARCB1 and LZTR1 mutant tumor cells. Interestingly, painful NOS CM also caused an increased response to the TRPA1 agonist cinnamaldehyde in our in vitro calcium imaging experiments. There is a growing consensus that TRPA1 is involved in mechanical hypersensitivity in different types of chronic pain under pathological conditions [14-16]. Among the cytokines elevated in painful NOS CM, CCL2 is a promising candidate for initiating and maintaining mechanical hypersensitivity. Previous studies showed that spinal administration of CCL2 in rats induced sustained painful mechanical hypersensitivity that lasted 4 days post-injection[17]. In addition, CCL2 acting on its receptor CCR2, sensitizes TRPA1, causing a sustained influx of calcium into cells[18]. Based on these previous findings it is possible that elevated CCL2 in painful NOS CM, activates or sensitizes TRPA1 and thereby induces an amplified and sustained mechanical hypersensitivity. IL-8 is another promising candidate mediator of lasting mechanical hypersensitivity. Injection of IL-8 into the footpad of the naïve rat is known to cause mechanical hyperalgesia [19]. IL-8 acts through the receptor CXCR1/CXCR2 during inflammation.

Previous studies demonstrated that IL-8 is involved in long lasting mechanical hypersensitivity that persists even after an inflammatory response has resolved [20], again consistent with our in vivo and in vitro findings on painful NOS CM. Lastly, another study demonstrated that intraplantar injection of CXCL5 into the rat paw caused a dose-dependent reduction in mechanical pain thresholds compared to vehicle control, which returned to baseline levels by 24 hours [21]. Based on these findings, targeting elevated cytokines may be an effective strategy to treat mechanically evoked pain induced by painful NOS SWN CM.

Painful LZTR1 mutant CM did not induce hypersensitivity to light touch compared to non-painful LZTR1 mutant CM. However, acute pain-related behaviors were greater in response to CM from nonpainful LZTR1 mut CM than that from control medium and were greater yet in response to painful LZTR CM. Painful LZTR1 mutant CM also evoked a larger acute pain response than either SMARCB1 or NOS CM (p=0.0004 and p=0.0003, respectively). This finding leads us to speculate that tumors with mutations in LZTR1 may have a different signature of secreted cytokines that increase acute pain. In fact, painful LZTR1 CM was the only sample that contained an elevated level of GM-CSF by ELISA and the highest level of CXCL-1 compared to the other painful CMs. The role of GM-CSF in pain and inflammation has been previously examined [22]. GM-CSF receptors are expressed in a subset of small/medium diameter neurons that also express TRPV1. In vivo, intraplantar injection of GM-CSF in mice caused favoring of the affected paw [22].

In conclusion, our findings demonstrate that painful SWN CM sensitizes mice to painful stimuli and that both the magnitude and duration of the pain hypersensitivity, as well as the cytokine/chemokine content of these CMs differ as a function of the SWN-associated gene mutation. These data, derived using an expanded cohort of painful and non-painful CM, validate our previous in vitro results. Additional testing of painful CM in vivo may lead to a better understanding of the etiology of SWN-related pain and guide the rational development of personalized therapies for pain in patients suffering from these disorders.

## Methods

### Tumor cells and conditioned media

Schwannomatosis-related tumors were collected from surgical cases from January 2014 through March 2021, occurring at Johns Hopkins School of Medicine. Informed, written consent was obtained prior to surgery. The Institutional Review Board (IRB) of the Johns Hopkins School of Medicine approved consent forms and study design. All research was performed in accordance with relevant guidelines/regulations and approved by the IRB. The diagnosis of SWN was confirmed by Johns Hopkins Pathology. Self-reported pain scores and mutation status were collected prior to surgery. Human Schwannomatosis cell lines were established from participant tumors as described in Ostrow et al 2015. Cell cultures were maintained in D10 media (DMEM, 10% FBS, 5% penicillin/streptomycin) with mitogens (2uM Forskolin). Media for testing was collected from these cultures at 48h intervals at 80% confluence Conditioned medium (CM) was centrifuged to remove cell debris and passed through a 0.22 micron filter. CM was frozen in aliquots and stored at −80C for future use.

### ELISA

Protein levels in CM were normalized to SERPINE1. The internal controls of the Human Cytokine XL Array (R&D systems) and SERPINE1 consistently demonstrated the mean pixel density across all CMs tested. Therefore, SERPINE1 was used as the internal control protein to normalize CM input prior to cytokine ELISA. Levels of SERPINE1 were tested in undiluted CMs to determine concentration using the R&D Quantikine total SERPINE1 ELISA kit. CMs were then diluted to a final concentration of 2ng of SERPINE1 prior to the ELISA assays. All ELISA experiments were performed according to a similar framework with reagent variation determined by their kit-specific instructions (R&D systems). In brief, CM samples and standards were pipetted into a microplate pre-coated with monoclonal specific antibodies, where target polypeptides were bound and further tagged with target-specific enzyme-linked polyclonal antibodies before proportional color analysis of added development substrate. Absorbance at 450nM was measured. Each sample was performed in triplicate. The amount of protein in each sample (pg/ml) was interpolated from a standard curve (Graphpad Prism 10).

### Mouse cohorts and experiments

All experimental procedures were approved by the Institutional Animal Care and Use Committee of Johns Hopkins University School of Medicine and were in accordance with the guidelines provided by the National Institute of Health and the International Association for the Study of Pain.

### Dissociated Mouse DRG cell cultures

Dorsal root ganglia (DRG) were harvested from 12 week old C57BL6J mice into Complete Saline Solution (CSS; NaCl 137mM, KCl 5.3mM, MgCl_2_-6H_2_O 1mM, Sorbitol 25mM, HEPES 10mM, CaCl_2_-2H_2_O 3mM). DRG were digested with TM Liberase (Trituration Moderate; 0.35U/mL in CSS, 50mM EDTA) at 37°C for 20 minutes in a rotating wheel hybridization oven, followed by TL Liberase (Trituration Low; 0.25U/mL in CSS, 50mM EDTA, 30U/mL Papain) at 37°C for 15 minutes by the same method. These were resuspended in complete DRG medium (DMEM/F12, 10%FBS, 1% glutamine, 5% penicillin/streptomycin) containing BSA (1.5mg/mL) and Trypsin Inhibitor (1.5mg/mL) before mechanical dissociation and passage through a 70um mesh filter. DRG were then spotted in 20uL TI/BSA/DMEM on poly-L-lysine/laminin coated coverslips and left to adhere for 1h in an incubator. Wells containing coverslips were then flooded with complete DRG medium.

### Calcium imaging

The day after establishment of primary mouse DRG cultures, culture medium was replaced with schwannomatosis cell CM and the cells were further incubated for 48 hours at 37 °C. Three separate DRG cultures were prepared for each CM to be tested. Four coverslips containing at least 30 neurons/slip were treated with CM. A minimum of 100 DRG neurons were tested per condition. Cells were treated with CM for 48 hours before loading with 2 μM fura-2 acetoxymethyl ester (Molecular Probes) in calcium imaging buffer (CIB, containing in mM: 130 NaCl, 3 KCl, 2.5 CaCl_2_, 0,6 MgCl_2_, 10 HEPES, 1.2 NaHCO_3_, 10 glucose, pH 7.45, 290 mOsm adjusted with mannitol). Coverslips with fura-2-loaded cells were mounted on an inverted fluorescence microscope (TE200, Nikon). Images were acquired with a cMOS camera (NEO, Andor) using an excitation filter wheel (Ludl) equipped with 340 and 380 nm filters. Data were acquired using NIS Elements imaging software (Nikon). Fluorescence changes are expressed as the ratio of fluorescence emission at 520 nm upon stimulation at 340 to that upon stimulation at 380 nm (F340/F380). Capsaicin experiments: Capsaicin was dissolved in CIB at a net concentration of 25 nM. During continual imaging at ∼2 sec intervals cells were perfused with 25 nM Capsaicin for 30 seconds with a recovery period of washing with unmodified CIB. Cinnamaldehyde experiments: Similarly, DRG cells were exposed to 200 uM cinnamaldehyde/CIB solution for 2 minutes followed by a 4 minute washout with CIB. The datapoints were corrected for differing baseline readings of Fura-2. Perfusion with KCl at a concentration of 50mM was used to identify viable neurons. Area under the curve was calculated for each treatment.

### Conditioned media Injections

Eight week old C57BL6J mice were placed in an immobilization cone and injected with 10uL of conditioned media in the right hind paw using a 20g insulin syringe. Ten mice were injected per conditioned media. Injections were repeated for 48 hours, 3 injections total. Mice were injected in groups of 20, (2 CMs; 10 mice per CM) to simplify handling and subsequent testing. CMs were deidentified and the experimenter was blinded to the code for the injections and subsequent behavioral assays to eliminate bias. All samples were the same in volume, color, and viscosity.

### Post-injection observations for acute pain behaviors

Immediately following day 1 injections all animals were recorded for a 10min observation period. Mice were placed into vertical plexiglass watch cylinders arranged in an arc of 5 opposite a mounted camera, in such a position to see all angles inside each cylinder. The mice were recorded from the placement of the first mouse in the group of five until 10 minutes after the placement of the last in the group. The observation period of each mouse was defined as 10 minutes, beginning when they were placed in the watch cylinder, and was used during the review and scoring of these recordings. The recorded 10min observation periods were then scored for reactive behavior (licking, flinching). Each instance of deliberate licking of the underside of the injected paw was considered a Lick, while each instance of rapid recoil or wave of the injected paw was considered a Flinch. Particular attention was paid to distinguish reactions from grooming behavior.

### Von Frey Assessment

Following the final injection, CM injected mice were placed into plexiglass cages in groups of 5 on top of a wire-mesh platform and given an acclimation period of twenty minutes, coinciding with the time intervals 1h, 24h, and 48h following final injections. Following this period, wire filaments of eight increasing gauge and force (0.02g, 0.04g, 0.07g, 0.16g, 0.4g, 0.6g, 1g) were applied to the hindpaws of each mouse to the point of bending six times per filament size. The response for each filament introduction to each paw was recorded as positive or negative. A dose-response frequency curve was calculated for each mouse.

### Data Analysis and Statistics

**(See also Supplementary Table 1)**

For ELISA: The level of specific cytokines were determined by interpolation of a standard curve using Graphpad Prism10. Data from Painful CM (n=12 samples) were grouped and compared to Non-painful CM (n=8 samples). Two-tailed unpaired t-tests with Welch’s correction were used to determine significance (Graphpad Prism 10). p value <0.05 was considered significant.

For calcium imaging: The area under the curve of fura-2 ratio, with subtraction of the baseline obtained prior to each stimulus, was used to compare the effects of painful vs. non-painful CM. Brown-Forsythe and Welch ANOVA tests with Dunnett’s T3 multiple comparisons test were used to compare the effects of CM groups (Graphpad Prism10). In addition, the percentage of cells responding to capsaicin, or cinnamaldehyde stimulation was analyzed. An individual cell’s response to stimuli was considered to be positive if the fura-2 ratio was greater than 0.1 after baseline subtraction. Chi-squared analysis was performed to test significance between the percentage of cells responding to painful and non-painful CM. p value <0.05 was considered significant.

For acute pain: Brown-Forsythe One-way ANOVA test and post-hoc Dunnett’s T3 multiple comparisons test were performed to determine significance between groups using Graphpad Prism 10.

For Von Frey analysis: Two-way ANOVA with repeated measures, and Tukey’s multiple comparison tests were performed to determine significance between groups using Graphpad Prism 10.

The authors will make materials, data and protocols available upon request.

## Acknowledgements

The Blaustein Pain Foundation, Pamela Mars Wright Foundation, and The Johns Hopkins Neurosurgery Pain Research Institute supported this work. We thank members of the Hoke and Caterina labs for helpful discussions.

## Author Contributions

R.R. conducted experiments, performed data analysis, and co-wrote the manuscript. K.L.O. conceived the project, designed the experiments, conducted experiments, analyzed the data, and co-wrote the manuscript. A.B. and M.J.C. facilitated experimental design, assisted in data analysis, and revised the manuscript.

## Additional information

Disclosure: None

