## Supplementary Figure for "Conditioned medium from painful schwannomatosis tumors increases pain behaviors in mice"

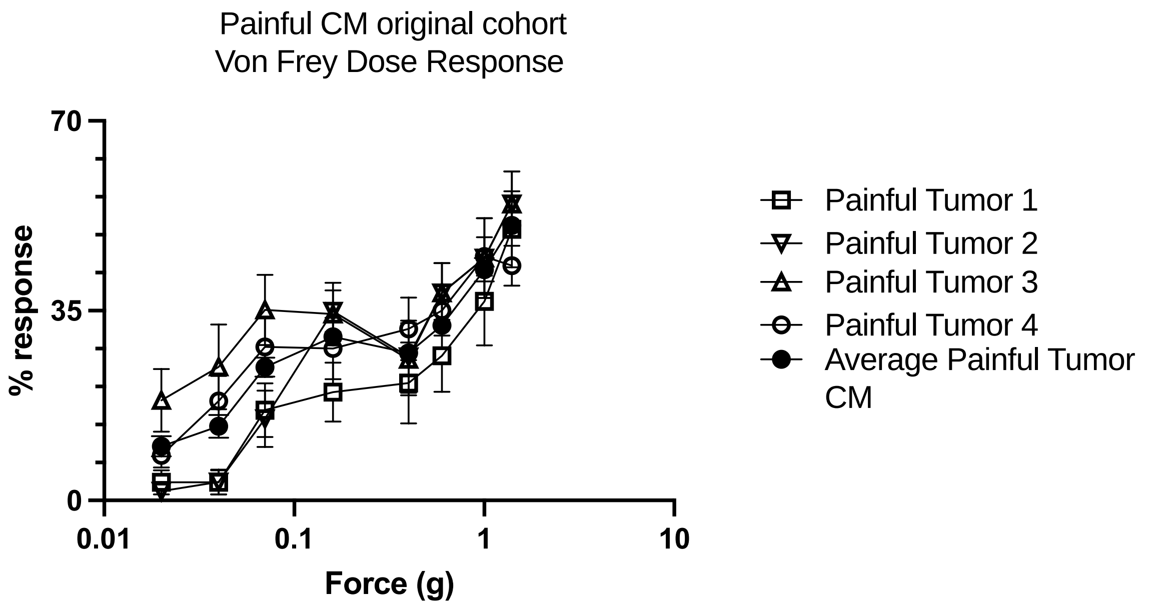


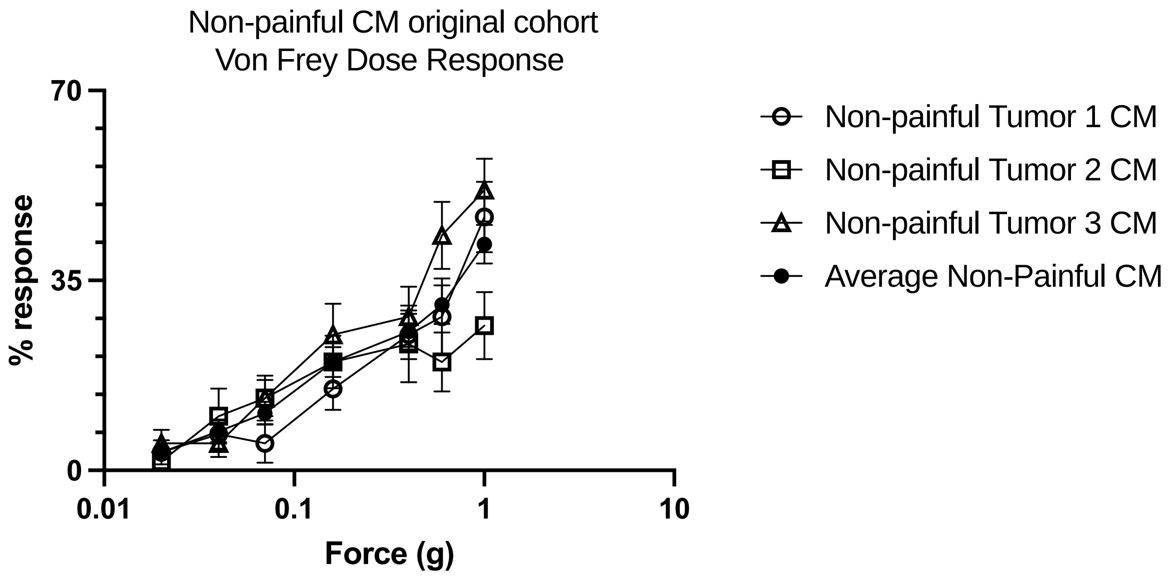


**Supplementary Figure 1:** Von Frey doses response for each painful (Top Panel) and non-painful (bottom panel) tumor CM

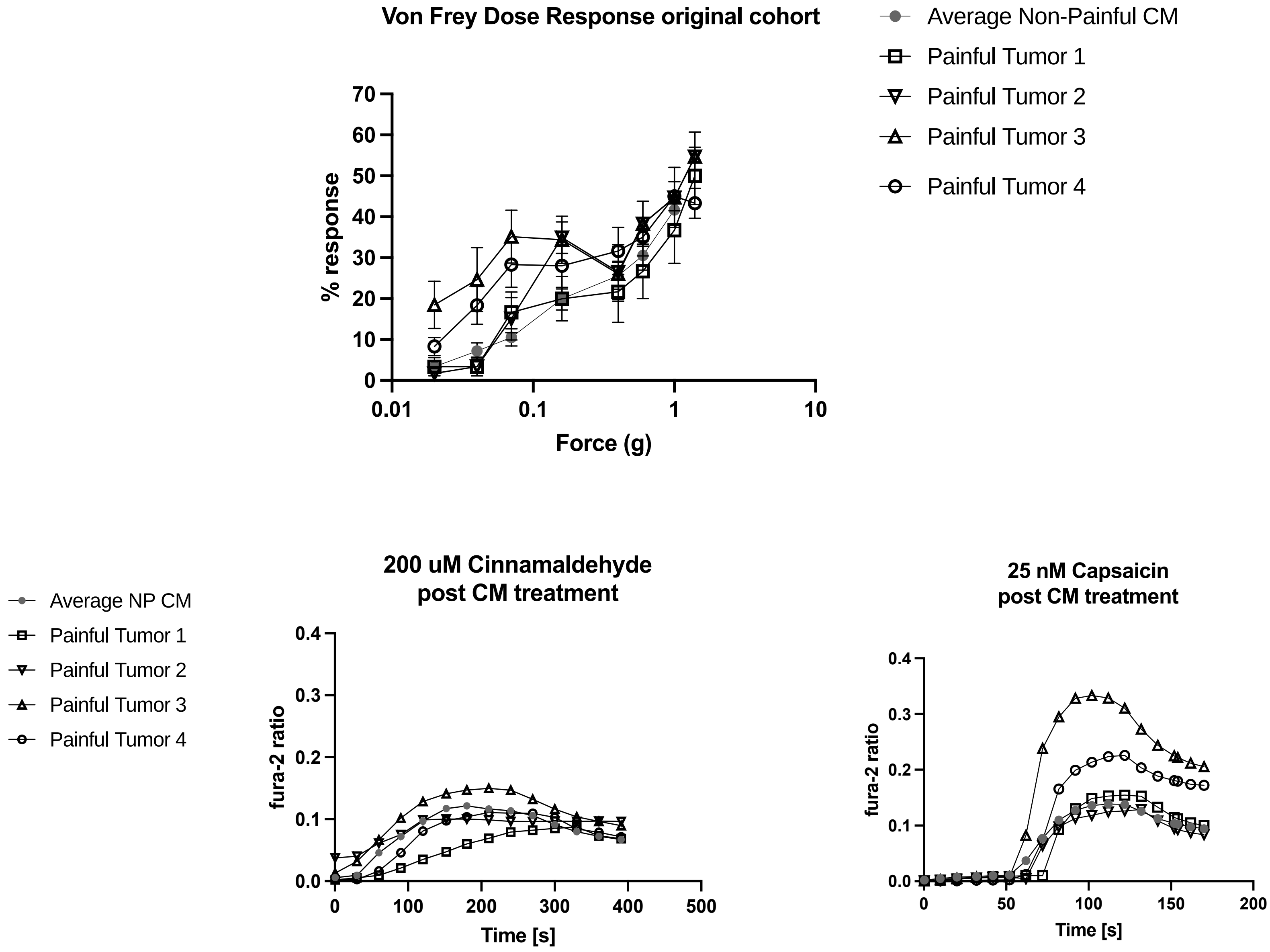


**Supplementary Figure 2:** Top panel: Von Frey doses response for each painful tumor CM and the average of non-painful tumor CM. Bottom panel: Corresponding calcium imaging time course DRG neurons pre-treated with CM.
